# A dual sgRNA library design to probe genetic modifiers using genome-wide CRISPRi screens

**DOI:** 10.1101/2023.01.22.525086

**Authors:** Alina Guna, Katharine R. Page, Joseph M. Replogle, Theodore K. Esantsi, Maxine L. Wang, Jonathan S. Weissman, Rebecca M. Voorhees

## Abstract

Mapping genetic interactions is essential for determining gene function and defining novel biological pathways. We report a simple to use CRISPR interference (CRISPRi) based platform, compatible with Fluorescence Activated Cell Sorting (FACS)-based reporter screens, to query epistatic relationships at scale. This is enabled by a flexible dual-sgRNA library design that allows for the simultaneous delivery and selection of a fixed sgRNA and a second randomized guide, comprised of a genome-wide library, with a single transduction. We use this approach to identify epistatic relationships for a defined biological pathway, showing both increased sensitivity and specificity than traditional growth screening approaches.

## Introduction

In higher eukaryotes, complex phenotypes are facilitated not only by genetic expansion, but by the combinatorial effects of genes working in concert (Badano & Katsanis, 2002; Hartman IV et al., 2001). Evolutionarily, this complexity affords both genetic redundancy and the ability to undergo rapid cellular adaptation, which ensures phenotypic robustness upon loss or mutation of any particular gene. Indeed, most fundamental processes are buffered by components with partially overlapping function including protein quality control (i.e. protein folding chaperones and E3 ubiquitin ligases), cellular stress response (i.e. the heat shock response and the ubiquitin-proteasome system), and protein biogenesis (i.e. targeting and insertion to the endoplasmic reticulum [ER]) (Morishima et al., 2008; Itakura et al., 2016; Rodina et al., 2016; Rutherford & Lindquist, 1998; Lehner et al., 2006). However, this creates technical challenges to genetically interrogating biological pathways and assigning gene function in mammalian cells. For example, loss of only ∼1/4 of the ∼10,000 genes expressed in a typical cell will result in any detectable growth phenotype (Winzeler et al., 1999; Costanzo et al., 2010; Tsherniak et al., 2017; Behan et al., 2019).

To address these challenges, genetic modifier screens have traditionally been a powerful tool for defining gene function, identifying missing components of known pathways, establishing disease mechanisms, and pinpointing new drug targets (Eshed et al., 2001; Eshed et al., 1999; Ding et al., 2016; Hannan et al., 2016; Ahmad et al., 2009; Hurd et al., 2013; Ham et al., 2008; Najm et al., 2018; Diehl et al., 2021; Wong et al., 2016). Forward genetic modifier screens rely on genetic ‘anchor points’ as a baseline for determining whether subsequent mutations, generally induced through random mutagenesis, result in buffering or synthetic phenotypes. In practice, this ‘anchor’ is established in a model organism or cell, often requiring extensive manipulation to generate a specific knockout in either organisms or cells, or isogenic mutant cell lines (Soldner et al., 2011; Perreault et al., 2005; Johnston, 2002). Apart from being technically cumbersome, classic forward approaches lack the ability to systematically assess genetic interactions on a genome-wide scale. The advent of CRISPR-based techniques has expanded this ability by allowing for (i) the generation of specific genetic perturbations in the form of knock-outs or knock-downs and (ii) the performance of unbiased genome-wide forward genetic screens to identify the genetic basis of an observed phenotype.

The majority of genetic modifier screens in human cells leverage a CRISPR cutting based approach (Zeng et al., 2019; Hickey et al., 2020; Kramer et al., 2018; Chai et al., 2020; DeWeirdt et al., 2020). However, Cas9-mediated DNA cutting is toxic to cells because it activates the DNA damage response, which is fundamentally problematic for genetic interaction analysis where multiple genomic sites are targeted (Fu et al., 2013; Hsu et al., 2014). Additionally, cells readily adapt and compensate for loss-of-function mutations over time, diminishing observed phenotypes when isogenic knockout cell lines are required (Norman et al., 2019). Moreover, relying on a genetic knock-out approach is often not amenable to the study of essential genes. A more acute strategy, CRISPR interference (CRISPRi), circumvents many of these issues and offers several advantages, notably the ability to create homogenous, titratable knock-down of genes without generating double-stranded DNA breaks (Doench, 2018). CRISPRi relies on a catalytically dead Cas9 (dCAS9) fused to a repressor domain, which, when guided by a sgRNA targeted to a particular promoter, results in the recruitment of endogenous modulators that lead to epigenetic modifications and subsequently gene knock down (Gilbert et al., 2013; Gilbert et al., 2014; Horlbeck et al., 2016; Qi et al., 2013).

We therefore envision that a strategy to query epistatic relationships acutely and systematically at scale, compatible with the sensitive phenotypic read-out afforded by a fluorescent reporter, would be a powerful tool for assigning genetic function. Towards this goal, we coupled existing CRISPRi technology with a simple and flexible dual-sgRNA library design that is compatible with multi-color FACS-based reporter screens. Our library design, which acutely delivers both a genetic ‘anchor point’ guide and a second randomized guide in a single plasmid, allows us to perform genetic modifier screens for essential and non-essential genes on a genome-wide scale. As a proof of principle, we applied this approach to dissecting the complex parallel pathways that mediate tail-anchored protein insertion into the endoplasmic reticulum (ER). This approach will be broadly applicable for (i) identifying functional redundancy, (ii) assigning factors to parallel or related biological pathways, and (iii) systematically reveal genetic interactions on a genome-wide scale for a given biological process.

## Results

### Dual sgRNA library design and construction

We developed a strategy to construct and deliver a library containing a fixed pre-determined guide, our genetic anchor point, with a second randomized CRISPRi guide from a single lentiviral backbone at scale (Figure 1A). The basis of our second guide is the CRISPRi-v2 library, a compact, validated 5 sgRNA/gene library targeting protein-coding genes in the human genome (Horlbeck et al., 2016). Ease of use was a primary focus of the library design which we addressed by (i) ensuring library construction relied on straightforward and inexpensive restriction enzyme cloning, (ii) developing a sequencing strategy that serves as a failsafe to ensure both guides are present, eliminating potential background, and (iii) designing the library such that the resulting data could be analyzed using an existing computational pipeline.

**Figure 1.**
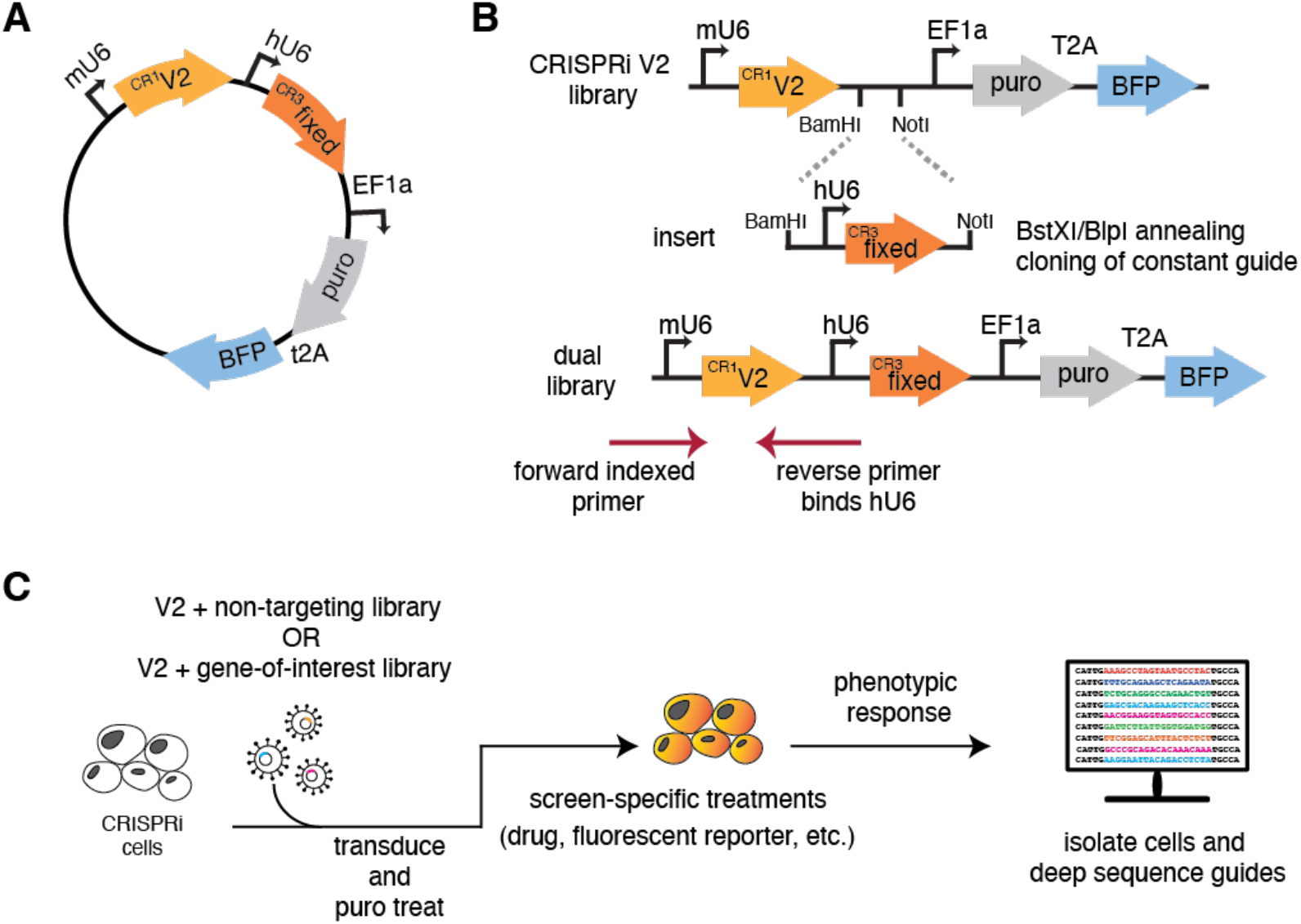
Dual-guide library design and construction. **(A)** Schematic of the dual sgRNA vector. Expression of the randomized CRISPRi-v2 sgRNA is driven by a mU6 promoter and the fixed guide is driven by a hU6 promoter, each flanked by unique guide constant regions (CR). Downstream, the EF1a promoter drives the expression of the puromycin resistance selectable marker and BFP. **(B)** Cloning a dual genome-wide library is comprised of two steps. First, a guide of interest is inserted using standard oligo annealing and ligation into a BstXI/BlpI cut backbone. Second, both CRISPRi-v2 library and the fixed guide are digested with complementary restriction sites (BamHI/NotI) and ligated at scale, resulting in an mU6- ‘V2 guide’-hU6-‘fixed guide’ library design. To sequence the resulting library, a standard 5’ indexed primer is coupled with a reverse primer that anneals to the hU6 region upstream of the inserted fixed guide. This strategy ensures only guides containing the fixed region are amplified for sequencing. **(C)** A general workflow for using our library design in any CRISPRi machinery containing cell.

A necessary requirement of our library design is identification of a pre-verified sgRNA that efficiently targets and depletes your gene of interest. This guide is first introduced by standard restriction enzyme cloning into a human U6 (hU6) and constant region 3 protospacer (CR3), hU6-CR3 cassette using sgRNA DNA oligos that can be inexpensively synthesized and purchased (Figure 1B). Using complementary restriction enzyme sites, the resulting hU6-CR3 cassette is ligated into the CRISPRi-v2 library at scale, resulting in an mU6-CR1-hU6-CR3 guide design (Replogle et al., 2020; Norman et al., 2019). As in the single element CRISPRi-v2 library, BFP and puromycin resistance genes are constitutively expressed, acting as fluorescent and selectable markers to identify guide containing cells.

Sequencing of the resulting library couples standard barcoded 5’ CRISPRi-v2 index primers with a new reverse primer complementary to the hU6 region, thereby only amplifying vectors containing the fixed sgRNA insert. This is important because during library construction, it is possible to produce a small fraction (we estimate <2%) that lack the fixed guide. Additionally, because this cloning strategy involves restriction enzyme digest of the CRISPRi-v2 library, there is loss of a small number of guides that contain these cut sites (∼1%, see Supplementary Table 1).

### Putative use of this dual sgRNA library for genetic modifier screening

To test this procedure, we first generated a library with a verified ‘non-targeting’ sequence as the fixed guide. Comparison with the standard CRISPRi-v2 library shows that we maintain similar guide coverage across the genome after accounting for expected loss of the restriction site containing guides (Figure S1A) (Horlbeck et al., 2016). The resulting sgRNA library allows for the acute knock-down of two separate targets without the need for additional selection markers, which simplifies both growth screens and the more sensitive fluorescent reporter-based flow cytometry screens. This design also removes the need to first make a cell line constitutively expressing a targeting or non-targeting sgRNA, thereby ensuring both the gene-of-interest and the genome-wide library are knocked down for the same period of time, diminishing the possibility of adaptation. Our library design is therefore compatible with a workflow that permits querying epistatic relationships with a variety of phenotypic readouts in any cells expressing the CRISPRi machinery (Figure 1C).

To test for genetic interactors at scale, one would conduct a screen using both the non-targeting library we have generated (available from Addgene, Library 197348), and a second library targeting a validated genetic ‘anchor point’ for your pathway of interest. Comparison of the results of these two screens, in the presence or absence of a characterized pathway component, will uncover and place factors in their respective pathway. We expect three possibilities. (i) Enhanced phenotypes in the ‘anchor point’ screen suggest synthetic effects, which would be indicative of factors in a parallel pathway. (ii) In contrast, diminished phenotypes in the anchor point screen would suggest factors in the same pathway. (iii) Finally, factors with phenotypes independent of our genetic ‘anchor point’ likely represent orthogonal genes.

### Developing a reporter assay to assess tail-anchored (TA) protein insertion at the endoplasmic reticulum (ER)

As a proof of principle, we tested the utility of our dual library by interrogating genetic interactors using a biological system known to contain at least two partially redundant pathways: tail-anchored membrane protein biogenesis. Tail anchored proteins (TAs) carry out essential functions including vesicle trafficking, organelle biogenesis, and cell-to-cell communication (Guna et al., 2022a). This family of integral membrane proteins are characterized by a single transmembrane domain (TMD) within 30-50 amino acids of their C terminus (Kutay et al., 1993). The proximity of the TMD to the stop codon necessitates that TAs be targeted and inserted into the membrane post-translationally. Though found in all cellular membranes, the majority of TAs are targeted to the ER using two parallel pathways: the Guided Entry of Tail-anchored protein (GET) and ER membrane complex (EMC) pathways (Stefanovic & Hegde, 2007; Schuldiner et al., 2008; Guna et al., 2018; Guna et al., 2022a).

In mammalian cells, the central components of the GET system are the targeting factor GET3, and the ER resident insertase composed of the heterooligomeric GET1/GET2 complex (Vilardi et al., 2011; Vilardi et al., 2014). The EMC pathway relies on targeting by the cytosolic chaperone, Calmodulin to the nine-subunit EMC insertase (Guna et al., 2018). The dependency of a particular TA on either set of factors is determined by hydrophobicity of its TMD, with more hydrophobic substrates relying on the GET, and those less hydrophobic relying on the EMC (Wang et al., 2010; Rao et al., 2016; Shao et al., 2017; Guna et al., 2018). However, substrates of intermediate hydrophobicity can utilize both pathways for targeting and insertion into the ER, potentially obscuring genetic relationships (Guna et al., 2018). We therefore reasoned that our dual-guide screening platform would be ideally suited to identify epistatic relationships between factors in these two pathways (Figure 2A).

**Figure 2.**
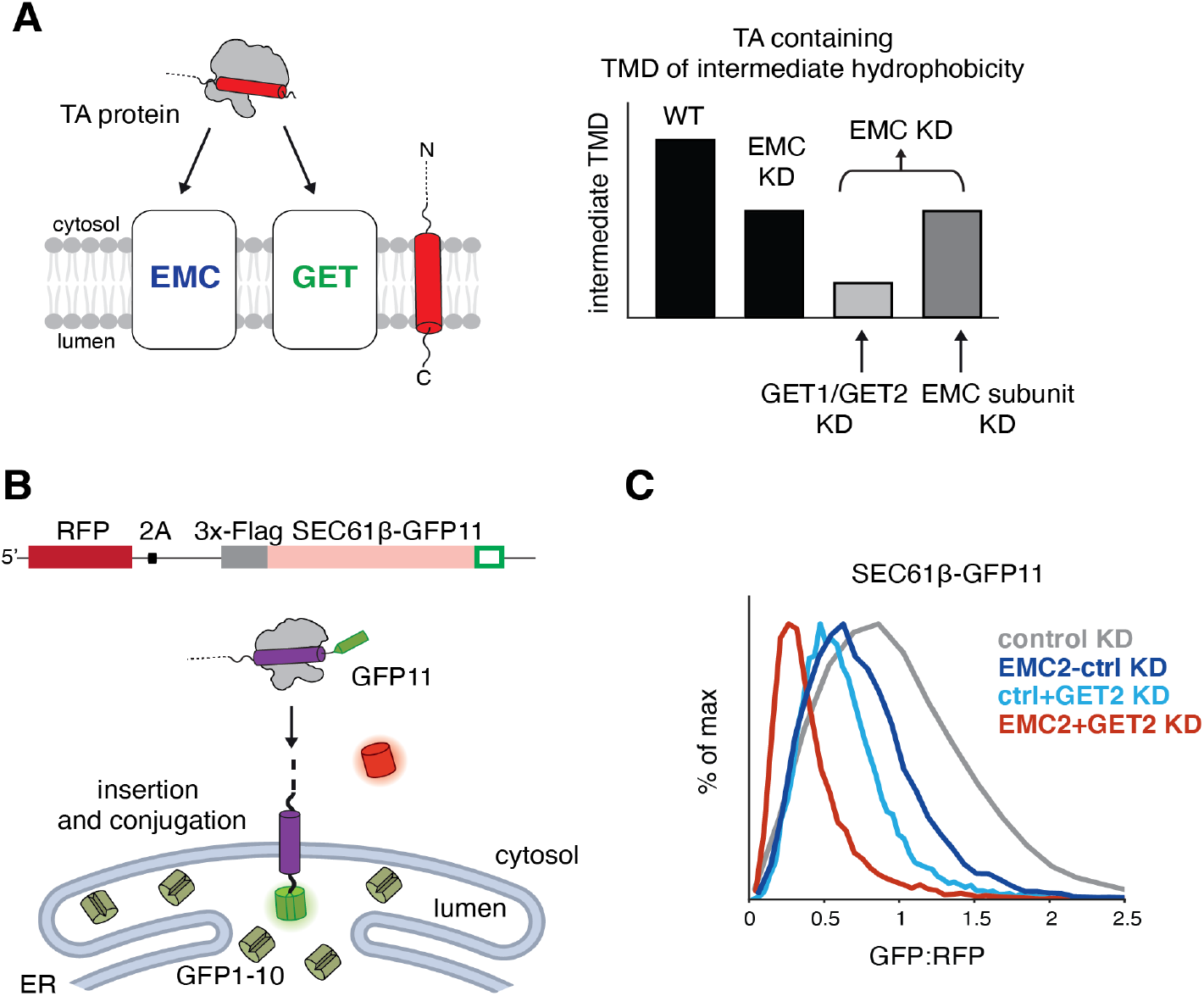
Querying tail-anchored (TA) protein biogenesis at the endoplasmic reticulum (ER). **(A)** (Left) TA proteins can be inserted into the lipid bilayer by either the EMC or GET insertases. (Right) TAs containing a moderately hydrophobic transmembrane domain such as SEC61β can use either EMC or GET1/GET2 to insert, obscuring strong effects on insertion when obstructing only one of these partially redundant pathways. Therefore, use of an EMC2 fixed guide dual library should uncover defined epistatic relationships between factors operating in either the GET or EMC pathways. **(B)** Schematic of the split GFP reporter system used to assess insertion of SEC61β into the ER. K562 cells expressing CRISPRi machinery were engineered to constitutively express GFP1-10 in the ER lumen. The 11^th^ β -strand of GFP is fused to the C-terminus of SEC61β, allowing for conjugation and fluorescence of the full GFP upon insertion into the ER membrane. RFP is expressed as a normalization marker, separated by a viral P2A sequence. **(C)** Depletion of EMC and GET pathway components in the SEC61β reporter cell line. The SEC61β cell line was separately transduced with dual guides targeting EMC2 alone, GET2 alone, EMC2 and GET2, or a non-targeting control. The GFP:RFP ratio, a measure of SEC61β insertion at the ER, is plotted for each dual guide.

To assess TA biogenesis using a FACS-based approach, we adapted a fluorescent split GFP reporter system to specifically query insertion into the ER (Figure 2B). For our reporter substrate, we chose SEC61β, which is an ER-localized TA that normally forms part of the heterotrimeric SEC61 complex (along with Sec61α, and γ) (Görlich et al., 1992; Hartmann et al., 1994; Görlich & Rapoport, 1993; Esnault et al., 1993). SEC61β contains a TMD of intermediate hydrophobicity and is known to use both the EMC and GET pathways for biogenesis (Guna et al., 2018). We constitutively expressed the first 10 β-strands of GFP (GFP1-10) in the ER lumen and appended the 11^th^ β -strand onto the C-terminal of the endogenous sequence of SEC61β (SEC61β-GFP11) (Inglis et al., 2020; Guna et al., 2022b). Successful insertion of SEC61β into the ER membrane would therefore result in complementation (GFP11 + GFP1-10) and GFP fluorescence. To generate cell lines compatible for screening, we engineered K562 cells to stably express ER GFP1-10 and the dCas9-KRAB(Kox1) CRISPRi machinery. Under an inducible promoter, we integrated the SEC61β-GFP11 reporter alongside a normalization marker (RFP) separated by a viral 2A sequence (Figure S1B). Expression of both the TA and RFP from the same open reading frame allows us to use the GFP:RFP ratio to identify factors involved in TA biogenesis while discriminating against those that have a non-specific effect on protein expression levels (i.e. transcription or translation).

### Interrogating TA insertion into the ER using dual sgRNA libraries

To permit screening with our dual-guide library design, we constructed a library using a previously validated EMC2 sgRNA as our ‘fixed’ guide (Figure S1C). EMC2 is a core, soluble subunit of the EMC complex, whose depletion leads to the post-translational degradation of the entire EMC via the ubiquitin-proteasome system (Volkmar et al., 2019; Pleiner et al., 2021).

Therefore, targeting EMC2 is sufficient to disrupt the EMC pathway for TA insertion, and therefore as our ‘genetic anchor’. Using our reporter line, we confirmed using programmed dual guides that loss of both the EMC complex and GET2 resulted in an additive effect of SEC61β insertion (Figure 2C). The enhanced effect of loss of GET2 in an EMC knockdown background validate the conceptual premise of our dual-guide screening approach at scale.

We therefore separately used both the EMC2 and a NT control library to transduce our K562 SEC61β reporter cell line, isolated cells that had perturbed GFP:RFP ratios by FACS, and identified the associated guides by deep sequencing. In parallel for comparison, we conducted a traditional growth screen with both the NT and EMC2 libraries in uninduced K562 SEC61β- GFP11 reporter cell lines (Figure S2A, Supplementary Table 2). As expected, in the NT-FACS screen loss of GET pathway components (GET2, GET3, and GET1) and all EMC subunits led to decreased SEC61β-GFP fluorescence, consistent with their established role in TA biogenesis. However, the EMC2-FACS screen showed markedly different results indicative of the genetic relationships between the EMC and GET pathway components (Figure 3A). First, when depleted on top of EMC2, the phenotype effects of loss of the main GET pathway factors is enhanced when compared to the NT screen. Second, the majority of guides targeting EMC subunits no longer have significant effects on SEC61β-GFP, consistent with their synergistic role with EMC2 (Volkmar et al., 2019; Pleiner et al., 2021). The exceptions are EMC2, likely because two guides targeting the same gene leads to a greater degree of knockdown, and EMC10, which has been suggested to have a separate regulatory role in TA biogenesis compared to the rest of the EMC complex (Coukos et al., 2021). Conversely, in both screens we also identified several novel ER-resident factors (RNF185, TMEM259 and FAF2) whose depletion leads to increased stability of SEC61β-GFP. Presumably, these putative quality control factors are responsible for recognizing and degrading over-expressed SEC61β from the membrane, but are agnostic to which biogenesis pathway was initially used for its insertion.

**Figure 3.**
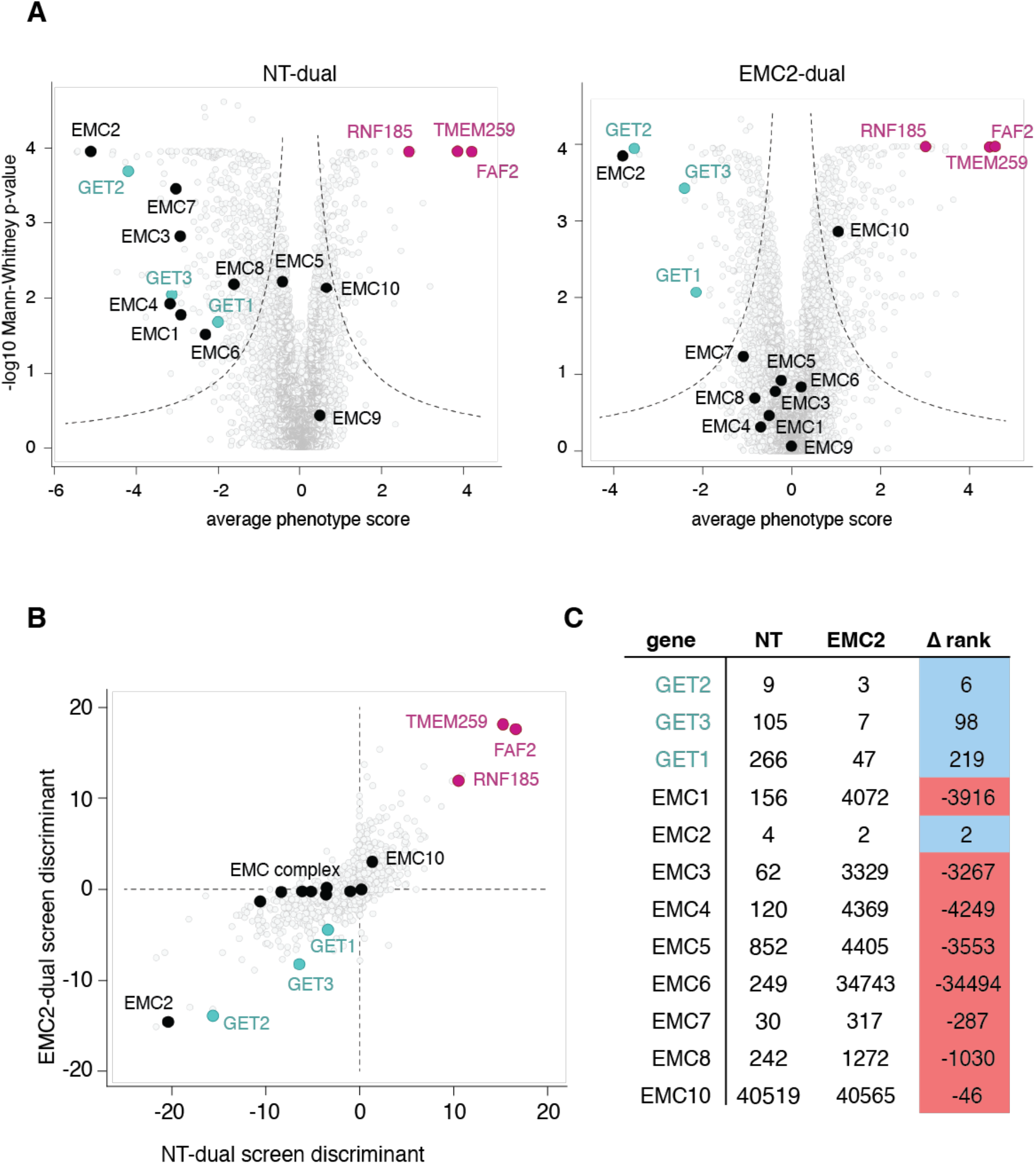
Dual-guide CRISPRi screen with SEC61β reveals genetic interactions between GET and EMC pathway components. **(A)** Volcano plot illustrating the phenotype for the three strongest guide RNAs versus Mann-Whitney p-values from two independent replicates of a genome-wide screen with either non-targeting dual (NT) or EMC2-dual libraries using the SEC61β -GFP11 reporter. Individual genes are displayed in gray, core factors of the GET pathway are highlighted in green, EMC subunits are highlighted in black, while putative stabilization factors are in pink. **(B)** A single discriminant score was computed for each gene in the screens investigating SEC61β -GFP11 stability, representative of the average phenotype score and significance of the hit in the respective screen. This metric allows direct comparison of both NT-dual and EMC2-dual screens. **(C)** Comparison of genes ranked by discriminant score in NT and EMC2-dual screens.

To facilitate the comparison of individual screens for identification of genetic interactors, we computed a discriminant rank as a single metric that combines both the phenotype score and the statistical significance of the effects of loss of each gene (Figure 3B). This allowed us to visualize the effects of a specific gene on SEC61β stability in the absence or presence of EMC2. Comparison of the NT- and EMC2-genome-wide FACS screens using the discriminant score highlighted the three broad categories of factors we anticipated: members of the GET pathway which show a synthetic effect with EMC, members of the EMC pathway which effectively ‘drop out’ in absence of EMC2, and factors which operate orthogonally from both pathways and are therefore unchanged in the two conditions (Figure 3C).

To confirm a subset of the observations predicted by our reporter-based screens, we conducted arrayed assays with programmed dual guides. Using our SEC61β -GFP11 reporter, we show that depletion of both EMC2 and GET3 has an enhanced effect on biogenesis compared to obstructing either pathway individually. This effect is likely specific to substrates of intermediate TMD hydrophobicity, as squalene synthase (SQS), a TA with known EMC dependency is only affected in the absence of EMC2 (Figure 4A) (Volkmar et al., 2019). Additionally, depletion of the putative quality control components RNF185, TMEM259 or FAF2 have affects the stability of SEC61β (Figure 4B), but not SQS or the GET substrate VAMP (Figure S3A). Indeed, RNF185 and TEMEM259 have been recently identified as members of a novel arm of ER-associated degradation (ERAD), while FAF2 has been previously associated with ERAD (van de Weijer et al., 2020; Xu et al., 2013; Lee et al., 2008).

**Figure 4.**
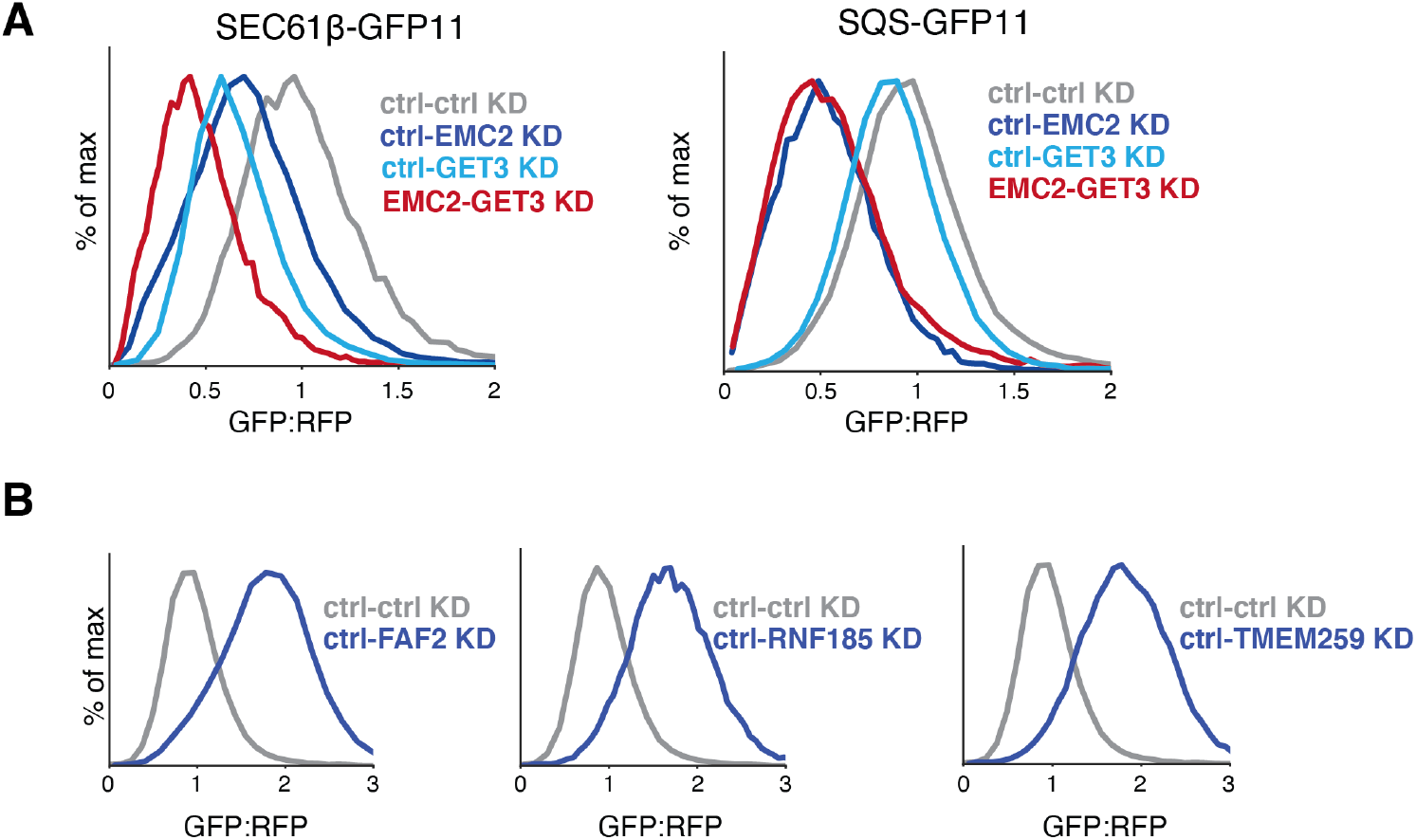
Validating effects of factors on TA biogenesis. **(A)** Integration of the TA proteins SEC61β -GFP11 or SQS-GFP11 into the ER was assessed in K562 cells that expressed the indicated programmed dual guides. GFP fluorescence is shown relative to a normalization marker (RFP) as determined by flow cytometry, and the results displayed as a histogram. **(B)** Biogenesis of SEC61β -GFP11 was assessed as in (A) with the presence of guides targeting the indicated genetic targets.

To illustrate the efficacy of our strategy, we compared the results of our FACS-based dual-guide library screen to a more traditional growth screening approach. Using growth as the metric, there is no increased genetic reliance on GET pathway components in the absence of EMC2 (Figure S2B, Supplementary Table 3). This is consistent with the observation that a substantial number of genes with transcriptional phenotypes have negligible growth phenotypes (Replogle et al., 2020). The significant number of hits both in the presence and absence of EMC2 are essential genes, occluding the possibility of detecting significant factors in the context of a particular biological pathway (Figure S2C). Given these results, hits identified from the growth screening approach would be particularly prone to off-pathway false positive and false negatives, necessitating substantial more follow-up to identify bona fide genetic interactors of the EMC. If we assume no previous knowledge of the relationship between the EMC and GET pathways, the growth-based approach clearly fails to identify genetic interactions that are crucial to elucidating its biological function. Thereby illustrating the efficacy and potential utility of our dual-guide screening approach.

## Discussion

We have developed a flexible, straightforward strategy to rapidly assess genetic interactions genome wide with high efficiency. Successful implementation of this approach does require sufficient prior knowledge of pathway or candidate gene of interest both to identify the fixed guide and design and validate an appropriate fluorescent reporter. However, the dual-guide strategy offers several practical advantages over existing genetic modifier screening strategies. Our approach eliminates the need to create and characterize a knock-out line for a particular gene of interest (Hickey et al., 2020; Feng et al., 2022; Westermann et al., 2022; Fu et al., 2013; Rossi et al., 2015). It also allows for the simultaneous delivery and selection of both targeted and genome-wide elements, resulting in less cell line construction and manipulation. The dual-guide library approach is compatible with multiple screening modalities while allowing for genome-wide perturbations, notably flow cytometry-based approaches where number of fluorophores may be limited. Finally, construction and use of new libraries is easy and rapid, with a two-step cloning process and reliance on existing sequencing and analysis pipelines. However, one minor caveat of the dual-guide system is that the addition of a second guide delivered on the same plasmid diminishes the efficiency of the fixed guide, but not by a significant amount (Figure S3B-C). This is evident in our system, with EMC2 coming out as a significant hit in the EMC2- dual-guide reporter screen. This can be ameliorated by the selection of a fixed guide that independently results in efficient knock-down, and the use of the recently described Zim3-Cas9 effector system, which has been shown to have stronger on-target knockdown compared to KOX1-Cas9 while maintaining minimal non-specific genome-wide effects (Alerasool et al., 2020; Replogle et al., 2022).

Recent studies have highlighted the success of FACS-based CRISPRi screens for the discovery of new factors (Guna et al., 2022b; Leto et al., 2019; Tsai et al., 2022; Morita et al., 2018). Our library extends the use case of this approach, for example allowing the study of processes that have parallel compensatory pathways, such as protein biosynthesis and degradation. Though a single screen with a programmed guide containing dual library is sufficient for most applications, performing an additional screen with a NT guide containing dual library provides additional data that could reveal critical genetic interactions. Another implication of our work is the relative paucity of information in traditional growth-based screens, with no additional perturbations.

Moving forward, we propose that additional up-front investment in developing a more targeted phenotypic read out, whether it be sensitivity to a compound or a reporter, is worthwhile when trying to establish genome-wide genetic interactions.

## Conclusion

The ability to genetically interrogate a biological process in mammalian cells on a genome-wide scale is a powerful tool to determine gene function. Here, we propose a simple advance to current CRISPRi sgRNA library construction that couples a genome-wide library with the simultaneous knock-down of a particular gene of interest. As a proof of principle, we use this design with a FACS-based reporter screen to show the relationships between the parallel pathways that mediate the insertion of TA proteins into the endoplasmic reticulum (ER). We not only faithfully reveal the known factors involved in this process, but can place them in either the GET or EMC pathways. We envision that these screening approaches represent a powerful strategy to unbiased and systematic identification of genetic interactors, capable of de-orphaning

## Materials and Methods

### Plasmids

Sequences used for in vivo analysis were derived from UniProtKB/Swiss-Prot and included: squalene synthase isoform 1 (SQS/FDFT1; **Q6IAX1**), vesicle associated membrane protein 2 (VAMP; **P51809-1**), and SEC61β (SEC61B, NP_006799.1). For expression in K562 cells, the transmembrane domain (TMD) and flanking regions of respective ER localized proteins were inserted into a backbone containing a UCOE-EF-1α promoter and a 3′ WPRE element (Addgene #135448) (Jost et al., 2017). The exception was the SEC61β construct used for the CRISPRi screens (RFP-P2A-Sec61b-GFP11) which was integrated into an SFFV-tet3G backbone (Jost et al., 2017). The GFP:RFP reporter system has previously been described (Chitwood et al., 2018) (Guna et al., 2018) and used in the context of CRISPRi screens (Guna et al., 2022b). The mCherry variant of RFP was used in all constructs, but is referred to as RFP throughout the text and figures for simplicity. For VAMP2, SQS and SEC61β, directly upstream of the TMD and flanking regions, the first 70 residues of the flexible cytosolic domain of SEC61β was inserted. Downstream, the GFP11 tag (RDHMVLHEYVNAAGIT) was inserted at the C-terminal separated by a 2-4X GS linker to allow for complementation with GFP1-10. In order to express GFP1-10 in the ER lumen, the human calreticulin signal sequence was appended preceding GFP1-10-KDEL as previously described (Cabantous et al., 2005) (Kamiyama et al., 2016) (Inglis et al., 2020).

Programmed dual sgRNA guide vectors were used to allow for the simultaneous depletion of genes (Replogle et al., 2020). Dual guide pairs included: EMC2-Control (GGAGTACGCGTCCGGGCCAA, GACGACTAGTTAGGCGTGTA), Control-GET2 (GACGACTAGTTAGGCGTGTA, GATGTTGGCCGCCGCTGCGA), EMC2-GET2 (GGAGTACGCGTCCGGGCCAA, GATGTTGGCCGCCGCTGCGA), Control-Control (GACGACTAGTTAGGCGTGTA, GACGACTAGTTAGGCGTGTA), Control-GET3 (GACGACTAGTTAGGCGTGTA, GGCTCCAGCGGCTCCACATC), EMC2-GET3 (GGAGTACGCGTCCGGGCCAA, GGCTCCAGCGGCTCCACATC), Control-FAF2 (GACGACTAGTTAGGCGTGTA, GCGGGTCAGGAGCGTAGAGG), Control-RNF185 (GACGACTAGTTAGGCGTGTA, GGCTGGCGTTAACTGTGCGG), Control-TMEM259 (GACGACTAGTTAGGCGTGTA, GCGGACGAGAAAGCGGAAGA). All reporter constructs and programmed dual guides are available upon request.

pCMV-VSV-G was a gift from Bob Weinberg (Addgene plasmid # 8454 ; http://n2t.net/addgene:8454 ; RRID:Addgene_8454).

### CRISPRi dual-guide library construction

Following selection and verification of a fixed guide, it is cloned into a hU6-CR3 cassette flanked by BamHI/NotI restriction cut sites (pJR152, Addgene 196280). The pJR152 backbone is compatible with standard BstXI/BlpI ligation with annealed oligos. Guide oligos must be ordered with custom overhangs (forward oligo: “ATG”-guide sequence-“GTTTCAGAGC”; reverse oligo: “TTAGCTCTGAAAC” – reverse complement of guide sequence – “CATGTTT”). For the NT and EMC2 libraries, the fixed guides were “GACGACTAGTTAGGCGTGTA” and “GGAGTACGCGTCCGGGCCAA”, respectively.

The two components of the dual-guide library are pJR152 containing the fixed guide of interest, and the CRISPRi-v2 library (Addgene Pooled Libraries #83969) (Horlbeck et al., 2016)(https://www.addgene.org/pooled-library/weissman-human-crispri-v2/). Construction of the dual-guide library essentially consists of restriction digesting both elements with BamHI/NotI and inserting the hU6-CR3-fixed guide element into the CRISPRi-v2 library at scale, resulting in an mU6-CR1-hU6-CR3 design previously described (Landisman & Connors, 2005).

Specifically, the pJR152 containing either the NT or EMC2 targeting guide was restriction digested at 37C for 3 hours with BamHI/NotI, and the resulting 400 bp gene fragment (containing the hU6-CR3-fixed guide element) gel purified. Approximately 30 ug of the CRISPRi-v2 top5 library was restriction digested with BamHI/NotI in the presence of shrimp alkaline phosphatase (rSAP) for 6 hours at 37C followed by heat inactivation at 65C for 5 minutes. Smaller amounts of the CRISPRi-v2 library can be digested, but a larger initial reaction will prevent repeat digestions and subsequent quality control checks. Ensure that no more than 10% of the reaction is enzyme to prevent star activity or inactivation of restriction enzymes. The resulting fragment of 8,800 base pairs was gel purified and eluted in a smaller volume.

Following recovery of both elements, either NT of EMC2 guide containing inserts were T4 ligated (ensure it is NEB #M0202M) with an insert to vector ratio of 1:2 for a 16 hours at 16C. Various vector:insert molar ratios were tested during piloting, with 1:2 resulting in the highest efficiency. A control ligation containing just the restriction digested CRISPRi-v2 library should be included.

To assess background, a small amount (0.5 ul of 20 ul) of the resulting ligations as well as the control were transformed into 10 ul of Stellar chemically competent cells (Takara #636763) using manufacturer guidelines. Various dilutions were plated (1/10^th^, 1/100^th^, and 1/1000^th^) with the resulting colonies counted on both control plates and dual library plates. Successful digestion and ligation should result in <2% background colonies, with the concern that single guides may pack much better than dual guides into lenti-virus, and therefore be over-represented.

To permit electrophoresis into MegaX cells at scale (ThermoFisher #C640003) the rest of the dual T4 library ligation for either NT or EMC2 dual libraries is selected with SPRISelect beads (Beckman Coulter B23317) and eluted in 20 ul of water. The entire resulting elution were electroporated into MegaX cells using manufacturer guidelines. Electroporated cells are allowed to recover and set up in an overnight culture of 200 ml LB supplemented with 100 µg/mL carbenicillin for each library. A smaller proportion of the culture was taken (1/1,000^th^ and 1/10,000^th^) and plated to allow for the estimate of resulting colonies and therefore guide coverage, with the expectation of maintaining 50X coverage for the 100,000 element CRISPRi- v2 library. Resulting NT and EMC2 dual libraries were amplified and barcoded by PCR using NEB Next Ultra ii Q5 MM (M0544L) and index primers and a unique reverse primer (CAAGCAGAAGACGGCATACGAGATggaatcatgggaaataggccctc) that binds in the hU6 region upstream of the fixed guide. The standard CRISPRi-v2 library was amplified in parallel to allow for the assessment of guide representation in dual libraries. (Horlbeck et al., 2016) SPRISelect beads (Beckman Coulter B23317) were used to purify the dual DNA libraries (349 bp), and purified DNA was sequenced using an Illumina HiSeq2500 with the same sequencing primer as the standard CRISPRi-v2 library (GTGTGTTTTGAGACTATAAGTATCCCTTGGAGAACCACCTTGTTG). The NT and EMC2 dual libraries are available on Addgene (Library 197348 and Library 197349, respectively).

### Cell culture and cell line construction

| **Cells** | **Source** |
| --- | --- |
| K562 KRAB-BFP-dCAS9 | Gilbert et al., 2014 |
| HEK 293T/17 | ATCC CRL-11268 |
| K562 KRAB-BFP-dCAS9 ER GFP1-10 | Guna et al., 2022 |
| K562-CRISPRi-Tet-ON-((ER)-GFP1-10)-(tet-RFP-P2A-SEC61β-GFP11) | This study, available upon request |

K562 cells expressing KRAB-BFP-dCas9 (Gilbert et al., 2014) were cultured in RPMI-1640 with 25 mM HEPES, 2.0 g/L NaHCO3, and 0.3 g/L L- glutamine supplemented with 10% FBS (or Tet System Approved FBS), 2 mM glutamine, 100 units/mL penicillin, and 100 μg/mL streptomycin. Cells were maintained between 0.25 × 10^6^ –1 × 10^6^ cells/mL. HEK293T/17 (ATCC CRL-11268) cells were cultured in DMEM supplemented with 100 units/mL penicillin and 100 μg/mL streptomycin. K562 and HEK293T cells were grown at 37C.

Cell lines expressing GFP1-10 in the ER lumen were generated as previously described (Inglis et al., 2020; Guna et al., 2022b). CRISPRi K562 cells were infected with lenti virus containing CalR(GFP1-10)-KDEL and sorted with a Sony Cell Sorter (SH800S) as single clones into 96- well plates. Clones were expanded and confirmed by complementation with a construct targeted to the ER appended to GFP11. To generate the SEC61β line used for screening line, lentivirus containing ER(GFP1-10) and RFP-P2A-SEC61β-GFP11 under an inducible promoter were co-infected at one copy per cell line in CRISPRi (expressing KRAB-BFP-dCas9) K562 Tet-ON cells (Gilbert et al., 2014). Cells were then single cell sorted, verified by induction with doxycycline (100 ng/ul), and confirmed to localize to the ER by microscopy. These cells are referred to as K562-CRISPRi-Tet-ON-((ER)-GFP1-10)-(tet-RFP-P2A-SEC61β-GFP11).

### Lentivirus production

Lentivirus was generated using standard protocols. Briefly, HEK293T cells were co-transfected with two packaging plasmids (pCMV-VSV-G and delta8.9, Addgene #8454) and either a desired transfer plasmid, or the dual libraries, using Transit-IT-293 transfection reagent (Mirus) (Stewart et al., 2003). Approximately 48 hours after transfection, the supernatant was collected and flash frozen. Virus was rapidly thawed at 37C prior to transfection.

### Flow cytometry reporter CRISPRi screens

CRISPRi screens were performed as previously described, with minor modifications (Gilbert et al., 2014; Horlbeck et al., 2016). Either the NT or EMC2 dual libraries were transduced in duplicate into 330 million K562-CRISPRi-Tet-ON-((ER)-GFP1-10)-(tet-RFP-P2A-Sec61b- GFP11) cells at a multiplicity of infection less than one. Throughout the screen, cells were maintained in 1L spinner flasks (Bellco, SKU: 1965-61010) at a volume of 1L. 48 hours after transfection, BFP positive cells were between 30-35%. At this point, cells began treatment with 1 µg/mL puromycin for three days to select for guide positive cells. Cells were given two days to recover after puromycin selection and the reporter was induced with doxycycline (100 ng/mL) for 24 hours and sorted on a FACSAria Fusion Cell Sorter. Cells were daily diluted to 0.5 × 10^6^ cells/mL to ensure that the culture was maintained at an average coverage of more than 1000 per sgRNA.

During sorting, cells were gated for BFP (to select only guide-positive cells) and RFP and GFP (indicating an expressing reporter). Cells were sorted based on the GFP:RFP ratio of the final gated population, and roughly 40 million cells with either the highest or lowest 30% GFP:RFP ratios were collected, pelleted, and flash-frozen. Genomic DNA of the cell pellets was extracted and purified using a Nucleospin Blood XL kit (Takara Bio, #740950.10). Guides were amplified and barcoded by PCR using NEB Next Ultra ii Q5 MM (M0544L) and index primers and a unique reverse primer (CAAGCAGAAGACGGCATACGAGATggaatcatgggaaataggccctc) that binds in the hU6 region upstream of the fixed guide. This ensures that only DNA containing both the v2 library and one of the fixed EMC2 or NT guides is amplified and sequenced. SPRISelect beads (Beckman Coulter B23317) were used to purify the DNA library (349 bp), and purified DNA was analyzed on an Agilent 2100 Bioanalyzer prior to sequencing using an Illumina HiSeq2500 using the same sequencing primer as the standard CRISPRi-v2 library (GTGTGTTTTGAGACTATAAGTATCCCTTGGAGAACCACCTTGTTG). Post-sequencing analysis was performed using the pipeline in https://github.com/mhorlbeck/ScreenProcessing (Horlbeck et al., 2016). Guides with fewer than 50 counts were excluded to ensure proper coverage. For each screen, the strongest 3 sgRNA phenotypes were used to calculate the phenotype score of each gene. The Mann-Whitney p-value was calculated using all 5 sgRNAs targeting the same gene compared to negative controls (Supplementary Table 2). Since screens were performed in biological duplicate, the sgRNA phenotypes were averaged. Discriminant scores were calculated as the product of the gene’s phenotype score and the Mann-Whitney p-value. Discriminant ranks for each screen were determined by ranking the list of genes from lowest to highest discriminant values, with the lowest score the highest rank.

### CRISPRi growth screens

To perform the growth screen, the same cells for flow cytometry screens infected with either NT or EMC2 dual libraries were harvested after recovery from puromycin selection as Day 0, and then again after 10 doublings on Day 18. 50 million cells from each biological duplicate and each library were harvested. Cells were maintained at an average coverage of more than 1000 per sgRNA during all points of the growth screen, and >99% BFP positive cells were confirmed at the time of harvesting. Resulting libraries were extracted, amplified, purified and sequenced identically as for the flow-cytometry based screen samples as described above (Supplementary Table 3).

### Flow cytometry

For all reporter assays, K562 CRISPRi cells containing ER(GFP1-10) were spinfected with lentivirus of indicated guides and knock down was allowed for 6 days. Cells were then spinfected with lentivirus containing the indicated reporters and analyzed by flow cytometry after 48-72 hours. All reporter experiments were performed in biological triplicate. All samples were either run on an NXT Flow Cytometer (ThermoFisher) or a MACSQuant VYB (Miltenyi Biotec). Flow cytometry data was analyzed either in FlowJo v10.8 Software (BD Life Sciences) or Python using the FlowCytometryTools package.

### Quantitative PCR

Quantitative PCR was used to analyze RNA levels after knockdown with dual or single guides. K562 cells expressing the CRISPRi machinery were infected with guides and after 8 days of knockdown, RNA was extracted and treated with DNaseI using a Direct-zol RNA MiniPrep Plus kit (R2072, Zymo). Purified RNA was reverse transcribed using the SuperScript III First-Strand Synthesis SuperMix for qRT-PCR kit (11752050, Invitrogen). Reactions were run on a StepOnePlus Real-Time PCR system and knockdown efficiency was calculated using the housekeeping gene HPRT1. Samples were collected and analyzed in triplicate with the means and standard deviations plotted. The primers used were: EMC2 (fwd AGACAGTTCCCTGGCAGTCAC, rev TCCACATTTTTCCCCTGGGCT); GET2 (fwd CCGGATCATGGGCTTTCACA, rev CCTGCTGGTCAGTTGTTCCT).

## Declarations

### Ethics approval and consent to participate

Not applicable.

### Consent for publication

Not applicable.

### Availability of Data and Materials

All materials necessary for dual-guide construction are available via Addgene, with accession numbers listed in the Materials and Methods. All programmed sgRNA, reporter constructs and cell lines are available from the corresponding author upon request. The sequencing datasets generated during the current study are available in the Caltech DATA repository and can be publicly accessed, https://doi.org/10.22002/3hvyj-yzq30

### Competing Interests

JMR consults for Maze Therapeutics and Waypoint Bio. JSW declares outside interest in 5 AM Venture, Amgen, Chroma Medicine, KSQ Therapeutics, Maze Therapeutics, Tenaya Therapeutics, Tessera Therapeutics, and Third Rock Ventures. RMV is a consultant and equity holder in Gate Bioscience. The Regents of the University of California with JSW as inventor have filed patent applications related to CRISPRi/a screening and Perturb-seq. JSW is an inventor on US Patent 11,254,933 related to CRISPRi/a screening. No potential conflicts of interests were declared by the other authors.

### Funding

Research reported in this publication was supported by: Howard Hughes Medical Institute (JSW), Center for Genome Editing and Recording 2RM1 HG009490-06 (JSW), Human Frontier Science Program 2019L/LT000858 (AG), the Heritage Medical Research Institute (RMV), the NIH’s National Institute of General Medical Sciences (DP2GM137412) (RMV), NIH F31 Ruth L. Kirchstein National Research Service Award NS115380 (JMR), Rosen Family fellowship (KRP), and Arie Jan Haagen-Smit Fellowship (KRP).

### Authors’ Contributions

K.P., A.G. J.R. and R.M.V. were responsible for conceptual design of the project. K.P., A.G., T.E., and M.W. performed experiments, K.P. and A.G. performed data analysis, and K.P., A.G., and R.M.V wrote the manuscript. R.M.V. and J.S.W. provided funding.

## Acknowledgements

We thank K. Hickey, R. Saunders, K. Popova and A. Inglis for helpful discussions. We thank the Whitehead Institute Flow Cytometry Core access to FACS machines and flow cytometers, and the Millard and Muriel Jacobs Genetics and Genomics Laboratory at Caltech for sequencing of screening libraries.

## Supplemental Figures and Figure Legends

**Figure S1.**
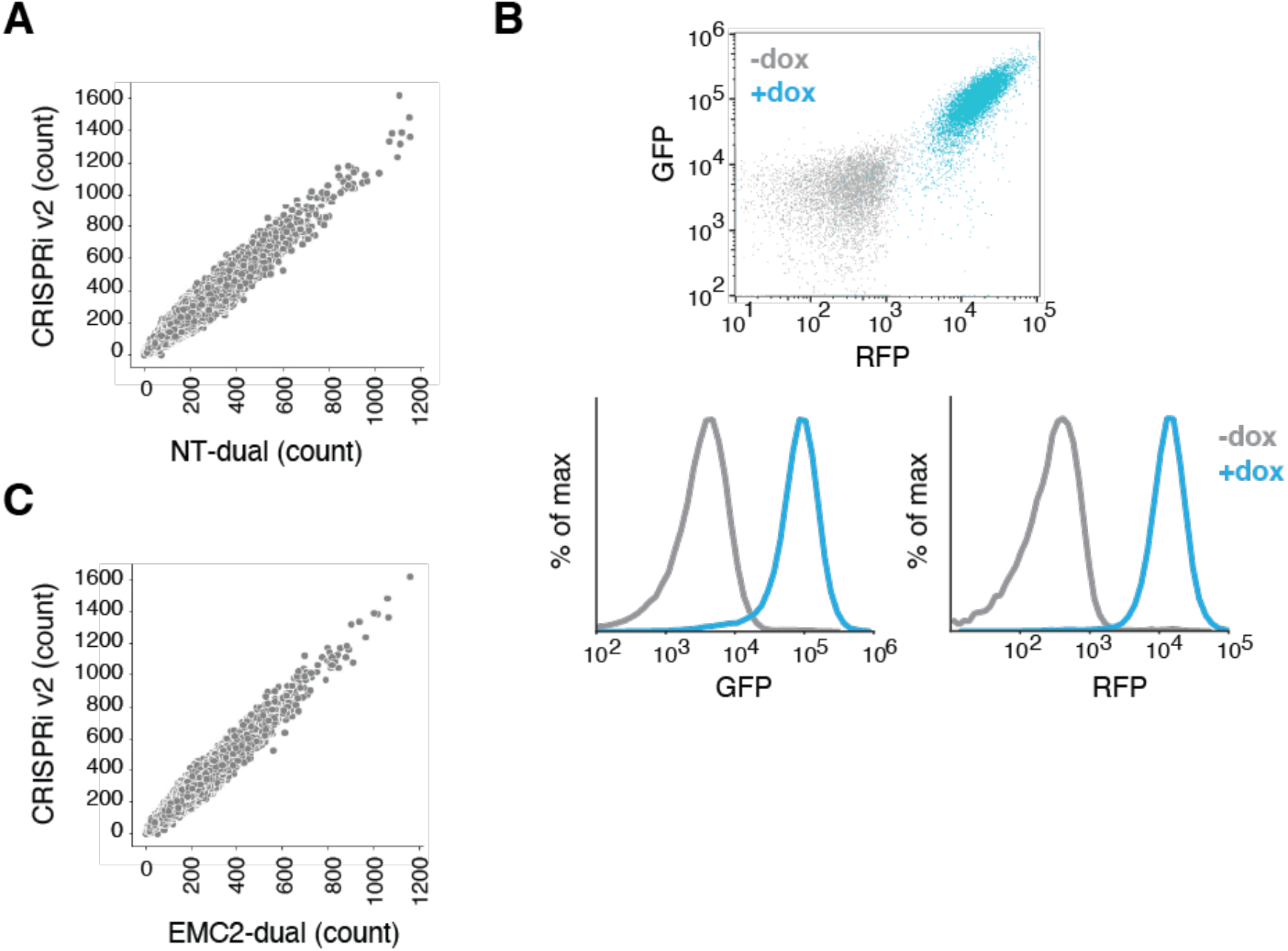
Dual library guide coverage and reporter line characterization. **(A)** Coverage of genome-wide guides in the NT dual library. Comparison of guide counts from the single CRISPRi-v2 library and the NT-dual library, after excluding guides which drop out due to restriction enzyme cutting during library construction. **(B)** K562 cells expressing GFP1-10 in the ER lumen and the SEC61β -GFP11 reporter under an inducible promoter are treated with doxycycline and analyze by flow cytometry. Green and red channels are shown separately. **(C)** As in (A) for the EMC2 dual library.

**Figure S2.**
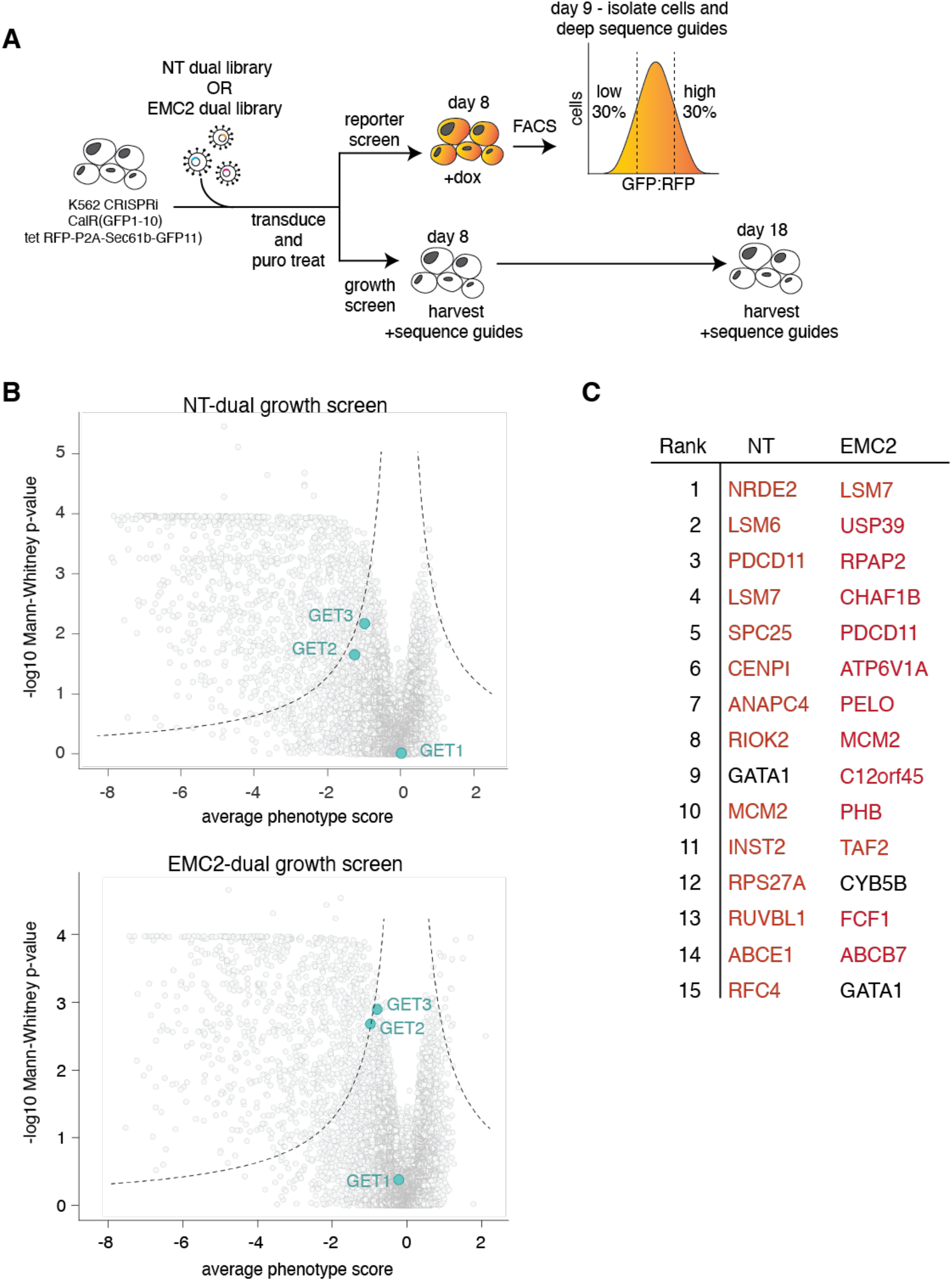
Growth screens with dual-guide libraries. **(A)** Schematic and timeline of CRISPRi fluorescent and growth screens with dual-guide libraries. **(B)** Volcano plots of growth screens for the three strongest guide RNAs versus Mann-Whitney p-values from two independent replicates of growth screens with the indicated libraries. Individual guides are displayed in gray, while core factors of the GET pathway are highlighted in pink. (**C**) Top ranked hits, as measured from discriminant scores, from (B), essential genes are highlighted in red (Tsherniak et al., 2017; Behan et al., 2019).

**Figure S3.**
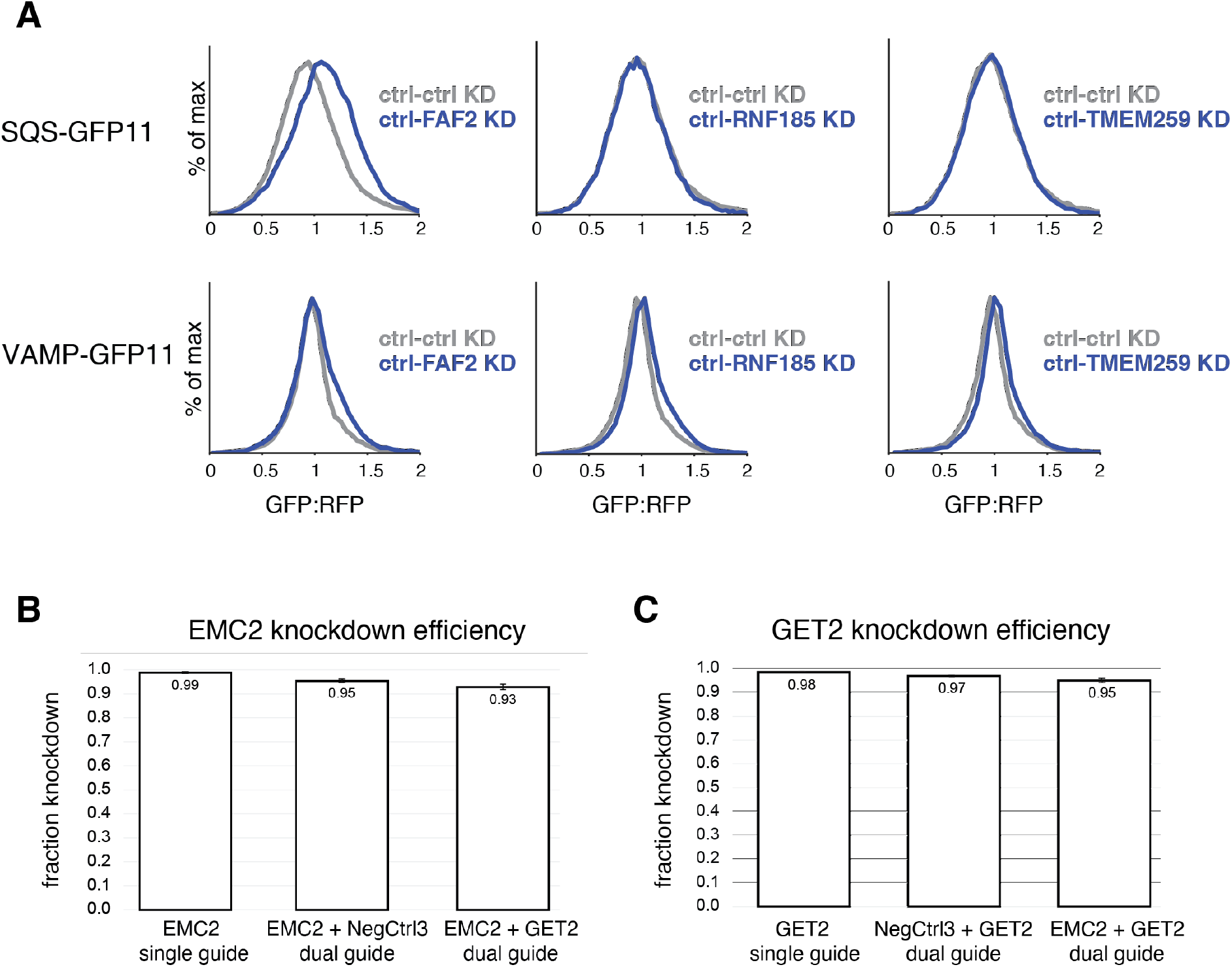
Investigating the specificity of putative TA quality control factors and assessing knock-down of dual-guides. (A) The stability of SQS-GFP11 and VAMP-GFP11 was assayed as in Figure 4B. (B) qPCR of EMC2 was assessed in K562 CRISPRi cells expressing the indicated guides to determine whether the context of a dual-guide affected knock-down efficiency (normalized to the housekeeping gene HPRT1). See methods for specific details. (C) As in (B) for the GET2 targeting guide.

## Supplementary File Legends

Supplemental Table 1. Restriction enzyme susceptible guides during dual library construction

Supplemental Table 2. Genome-wide FACS screens with non-targeting and EMC2 dual libraries for TA insertion at the ER

Supplemental Table 3. Genome-wide growth screens with non-targeting and EMC2 dual libraries for TA insertion at the ER

## Notes

### Summary of Updates

Supplemental figure S3B-C and additional references in the introduction have been added.

